# Monkeypox virus quadrivalent mRNA vaccine induces antibody responses and cellular immunity and protects mice against Vaccinia virus

**DOI:** 10.1101/2022.11.22.517500

**Authors:** Ye Sang, Zhen Zhang, Fan Liu, Haitao Lu, Changxiao Yu, Huisheng Sun, Jinrong Long, Yiming Cao, Jierui Mai, Xin Wang, Jiaxin Fang, Youchun Wang, Weijin Huang, Jing Yang, Shengqi Wang

## Abstract

There is an urgent need for efficient and safe vaccines against the monkeypox virus (MPXV) in response to the rapidly spreading monkeypox epidemic. In the age of COVID-19, mRNA vaccines have been highly successful and emerged as platforms enabling rapid development and large-scale preparation. Here, we have developed two MPXV quadrivalent mRNA vaccines, named mRNA-A-LNP and mRNA-B-LNP, based on two IMVs (A29L and M1R) and two EEVs (A35R and B6R). By administering mRNA-A-LNP and mRNA-B-LNP intramuscularly twice, mice have induced MPXV-specific IgG antibodies and potent Vaccinia virus (VACV)-specific neutralizing antibodies. Additionally, it elicited durable MPXV-specific killer memory T-cell immunity as well as memory B-cell immunity in mice. Furthermore, the passive transfer of sera from mRNA-A-LNP and mRNA-B-LNP-immunized mice protected nude mice against the VACV challenge. In addition, two doses of mRNA-A-LNP and mRNA-B-LNP were also protective against the VACV challenge in mice. Overall, our results demonstrated that mRNA-A-LNP and mRNA-B-LNP appear to be safe and effective vaccine candidates against monkeypox epidemics, as well as against outbreaks caused by other orthopoxviruses, including the smallpox virus.

## Introduction

Monkeypox, caused by the monkeypox virus (MPXV), is a zoonotic viral disease. Since May 2022, there have been more than 78,000 confirmed MPXV cases and more than 40 deaths worldwide, and transmission has occurred in 110 countries. Monkeypox has been declared a public health emergency by the World Health Organization. According to phylogenetic analysis, MPXV can be divided into two distinct clades, Central Africa (also known as the Congo Basin) and West Africa, with a high degree of sequence similarity between the two clades.^1^ Among them, The Central African clade tended to be more virulent, with a mean mortality rate of 10.6 % compared to 3.6 % for the West African clade.^2^ This may be because several open reading frames encoding immune evasion genes are disrupted in the West African clade, resulting in lower virulence in this clade.^3^ Nevertheless, it is the West African clade that caused the cases that are circulating in this outbreak.^4^

As a family of double-stranded DNA viruses, orthopoxviruses are composed of various viruses, including the Variola virus, Vaccinia virus (VACV), MPXV, and Cowpox virus.^5^ Orthopoxvirus infection or immunization confers immunity against other viruses in the genus.^6^ As a result, two vaccines have been approved by the FDA for the prevention of MPXV: ACAM2000, a second-generation live VACV vaccine, and JYNNEOS, an attenuated third-generation vaccine based on Modified Vaccinia Ankara (MVA). It has been reported that these two VACV-based smallpox vaccines are cross-protective against MPXV.^7^ However, smallpox vaccination does not completely protect against MPXV during the current outbreak, according to a recent study. ^8^ Additionally, ACAM2000 was a highly reproducing VACV vaccine that caused serious adverse events.^9^ These events include autoinoculation of the eye, generalized vaccinia, eczema vaccinatum, progressive vaccinia, myocarditis, and death.^10^ On the other hand, JYNNEOS, a replication-deficient vaccine, is expected to have the lowest incidence of severe adverse events.^9^ However, as live virus vaccines, JYNNEOS and ACAM2000 have undefined immune targets, and the impact of their gene products on immunity and adverse reactions remains unclear. Therefore, it is imperative to develop a vaccine specifically for MPXV to ensure complete protection.

In this study, the MPXV quadrivalent mRNA vaccines, mRNA-A-LNP and mRNA-B-LNP, were prepared using A29L, A35R, M1R, and B6R as antigenic targets. Orthopoxviruses can infect cells through two different mechanisms depending on whether they are intracellular mature viruses (IMV) or extracellular enveloped viruses (EEV).^11^ There are two IMV-specific proteins (A29L and M1R) in the quadrivalent mRNA vaccine, and two EEV-specific proteins (A35R and B6R). Among the orthologous homologous to VACV, the L1R and A27L are known neutralizing antibody targets for IMV; the B5R are known neutralizing antibody targets for EEV; however, the A33R is a target of complement-mediated cytolysis.^12, 13^ The combination of IMV- and EEV-specific immunogens have been found to provide more protection than either immunogen alone.^14^ It was reported that a DNA vaccine expressing A27L, A33R, L1R and B5R provided protection against lethal monkeypox,^12^ rabbitpox^15^ and vaccinia^16^ virus challenges. In contrast, vaccination with a single L1R provided a degree of protection against the lethal MPXV challenge but not against severe disease.^12^

Since the outbreak of COVID-19, mRNA vaccines have received unprecedented attention. There is a better immune response and safety associated with mRNA vaccines than with live virus and DNA vaccines.^17, 18^ Furthermore, as opposed to multiple punctures (scarification) percutaneous administration of live VACV vaccines^6^ and electroporation of DNA vaccines^15^, intramuscular administration of mRNA vaccines has reduced healthcare worker training and healthcare costs. So far, successful mRNA immunization resulting in protection from orthopoxviruses has never been reported. Here, we targeted MPXV homologs rather than VACV as antigenic targets to prepare MPXV quadrivalent mRNA vaccine, which induced MPXV-specific IMV- and EEV-antigen-specific IgG and potent VACV live-virus neutralizing antibodies. Meanwhile, immunized mice elicited a durable cellular response. Furthermore, mRNA-A-LNP and mRNA-B-LNP protected mice from VACV challenge.

## Results

### *In vitro* characterization of mRNA-A-LNP and mRNA-B-LNP

In this study, we developed potent MPXV quadrivalent mRNA vaccines that target two IMV-antigens and two EEV-antigens. Here, A29L, A35R, M1R and B6R, respectively, were chosen as antigenic targets for the mRNA coding sequences (Fig. 1a). The modified mRNA molecule begins with a 5’-cap 1, adheres to the 5’- and 3’-untranslated regions (UTRs), and then ends with a Poly-A tail. First, based on different mRNA design platforms, we constructed two groups of mRNA sequences, mRNA-A-mix and mRNA-B-mix. All mRNAs have been authenticated by capillary electrophoresis using the Agilent 2100 Bioanalyzer System (Supplementary information, Fig. S1). Based on Western blot analysis, both mRNA-mix detected bands at positions corresponding to each of the four antigenic proteins. It indicated that both mRNA-A-mix and mRNA-B-mix could successfully express the proteins (Fig. 1b). Afterwards, the mixed mRNAs were processed into LNP formulations. The resulting two quadrivalent mRNA vaccines were named mRNA-A-LNP and mRNA-B-LNP, respectively (Fig. 1c). The final stored mRNA-A-LNP and mRNA-B-LNP showed average particle sizes of 98.16 nm and 89.11 nm, respectively, and potentials of 5.3 mV and 6.9 mV. Meanwhile, over 90% of encapsulation rates were achieved by both (Fig. 1d). The cryo-transmission electron microscopy (TEM) analysis also revealed that both mRNA-A-LNP and mRNA-B-LNP particles exhibited a homogeneous solid spherical morphology, which indicated that the morphology of the LNPs loaded with the four mRNAs remained stable (Fig. 1e). According to these results, mRNA-A-LNP and mRNA-B-LNP can efficiently express MPXV proteins A29L, A35R, M1R, and B6R *in vitro*.

**Figure 1.**
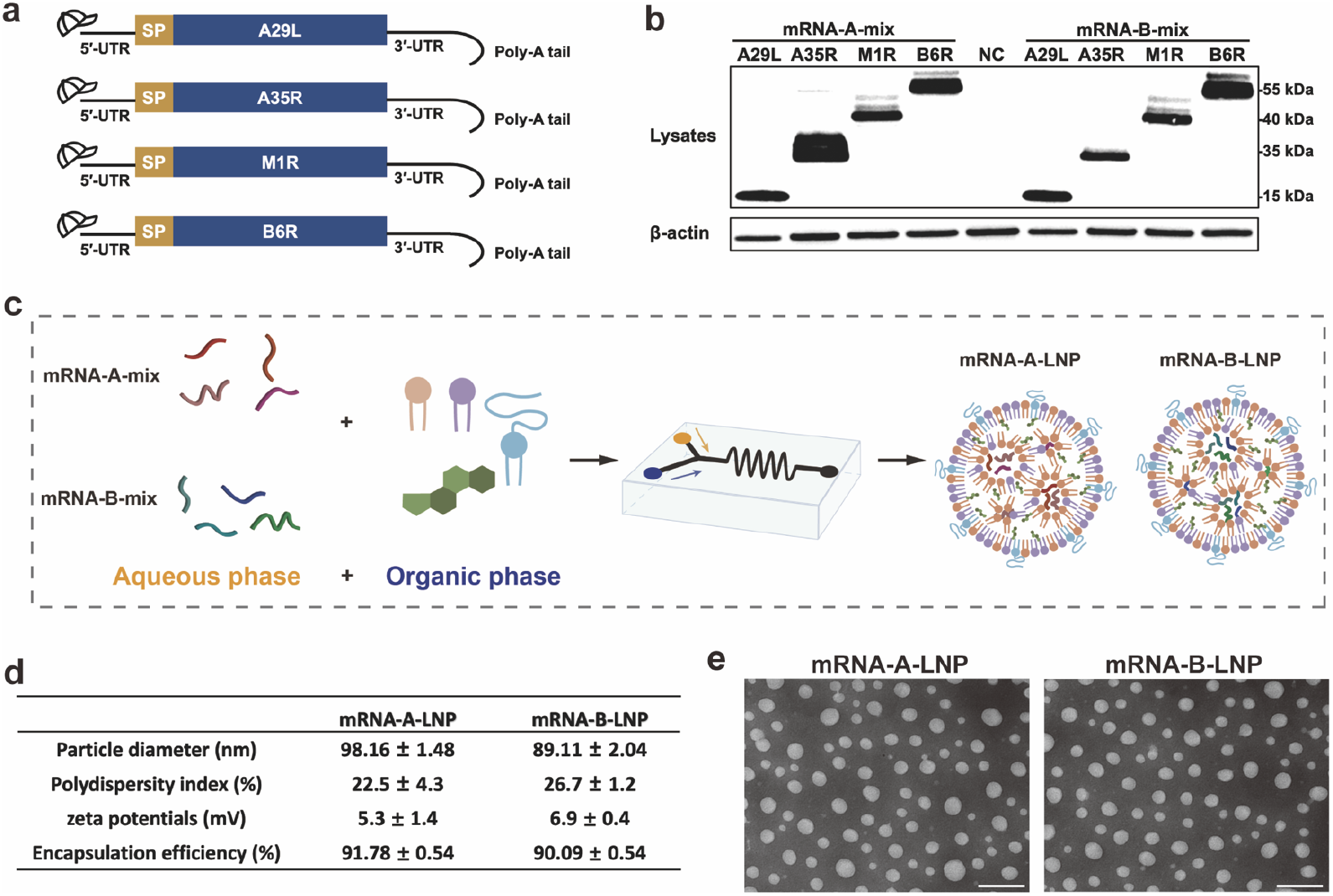
Design and encapsulation of mRNA-A-LNP and mRNA-B-LNP. (a) The mRNA construct expressing the MPXV-specific antigen A29L, A35R, M1R, B6R. (b) The MPXV-specific antigen A29L, A35R, M1R and B6R was expressed by mRNA in HEK293T cells. Cells were transfected with four mRNAs (1 μg/mL) each from mRNA-A-mix and mRNA-B-mix for 20 hours using Lipofectamine 3000 transfection reagent. (c) Preparation mechanism of mRNA-A-LNP and mRNA-B-LNP. Briefly, the four mRNAs were mixed in an acidic aqueous solution, then injected with organic phase lipids, and the mixture was extruded through a microfluidic chip. (d) The physicochemical parameters of mRNA-A-LNP and mRNA-B-LNP. Data are shown as mean ± SEM. (e) A representative transmission electronic microscopic (TEM) image presented the morphology of mRNA-A-LNP and mRNA-B-LNP. Scale bar = 200 nm.

### The prophylactic administration of mRNA-A-LNP and mRNA-B-LNP induces potent humoral immunity in mice

The immunogenicity and efficacy of mRNA-A-LNP and mRNA-B-LNP were also assessed in mice. Initially, female BALB/c mice were divided into three groups (n = 5) and immunized at intervals of 14 days (Fig. 2a). The mice were immunized with 40 μg of mRNA-A-LNP or mRNA-B-LNP, respectively, by twice intramuscular administration; naive mice served as the control group.

**Figure 2.**
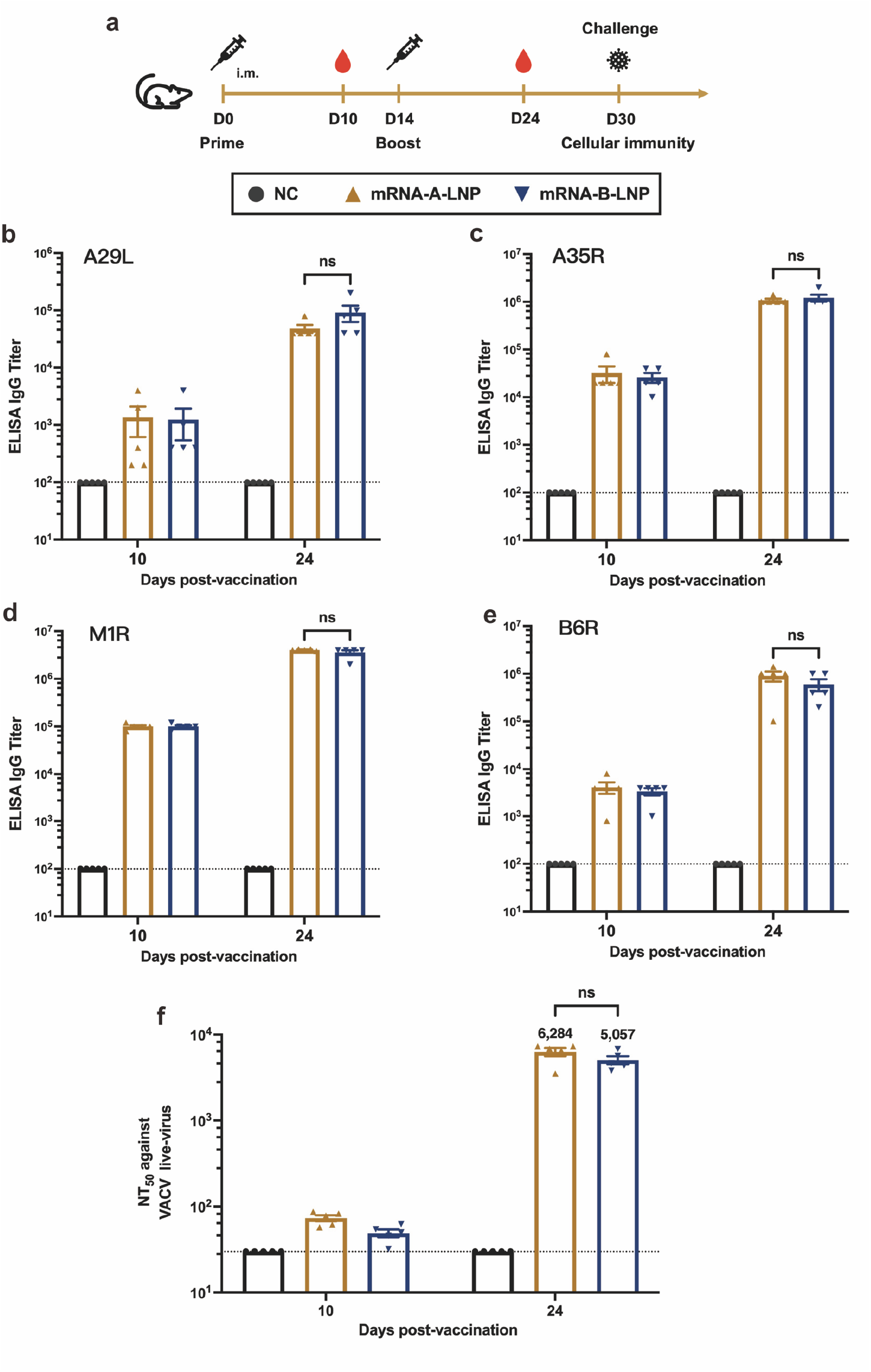
Humoral Immune Response in mRNA-A-LNP and mRNA-B-LNP-Vaccinated Mice. Female BALB/c mice were immunized with 40 μg mRNA-A-LNP vaccine (n = 5) or 40 μg mRNA-B-LNP vaccine (n = 5) or were designated negative controls (n = 5). Two intramuscular immunizations were on day 0 and day 14, respectively. Serum was collected 10 and 24 days after the initial vaccination. (a) Schematic diagram of immunization, sample collection, and challenge schedule. (b-e) The MPXV-specific antigen A29L, A35R, M1R and B6R IgG antibody titer was determined by ELISA. (f) The NT50 was determined by neutralizing antibody assay based on live VACV. The dashed line indicates the limit of detection of the assay. Data are shown as mean ± SEM. Significance was calculated using two-way ANOVA with multiple comparison tests (n.s., not significant; **p < 0.01).

Then, sera were collected from all mice 10 and 24 days after their first immunization to assess humoral immunity. Furthermore, no local skin reactions or inflammation were observed at the injection site. This suggests, unlike ACAM2000 (the damage at the vaccine site is often used as a marker of successful vaccination in highly replicating vaccines like ACAM2000), our vaccine does not undergo a significant cutaneous reaction, also known as “take”, which means no risks of autoinoculation or accidental vaccination.^6^ The binding antibody responses against MPXV antigens (A29L, A35R, M1R and B6R) were assessed by enzyme-linked immunosorbent assay (ELISA) 10 and 24 days after vaccination. Antibodies against all four target antigens were detected in all vaccinated mice and increased significantly over immunization frequency. The IgG titers against the four MPXV antigens at day 24 were 48,000 (A29L), 1,080,000 (A35R), 4,040,000 (M1R), 900,000 (B6R) for mRNA-A-LNP and 92,000 (A29L), 1,200,000 (A35R), 3,600,000 (M1R), 600,000 (B6R) for mRNA-B-LNP, respectively (Fig. 2b-e). The results indicated that mRNA-A-LNP and mRNA-B-LNP induce robust IgG against MPXV antigens.

The neutralizing antibody responses were assessed by live-virus neutralization tests using the VACV Tian Tan strain. After initial vaccination, neutralizing antibody titers were slightly above the detection limit, with median values of 73 and 49 for mRNA-A-LNP and mRNA-B-LNP, respectively (Fig. 2f). After booster immunization, neutralizing antibody titers increased by nearly 2 log10, with median values of 6,284 and 5,057 for mRNA-A-LNP and mRNA-B-LNP, respectively (Fig. 2f). Based on the MPXV antigen-specific ELISA and VACV live-virus neutralization tests, there was no statistical difference in the humoral response to mRNA-A-LNP or mRNA-B-LNP immunizations. In conclusion, these results suggest that the quadrivalent mRNA vaccine induced neutralizing antibodies against VACV and IgG antibodies against MPXV.

### The mRNA-A-LNP and mRNA-B-LNP induce long-term cellular immunity in mice

MPXV infection is primarily controlled by antibodies, but memory B cells and memory T cells play a role in the development of the humoral response to the monkeypox vaccine.^19^ Here, through the measurement of MPXV-specific germinal center (GC) B cells, follicular helper T (Tfh) cells, as well as CD4^+^ and CD8^+^ effector memory T (Tem) cells, we assessed the ability of mRNA-A-LNP and mRNA-B-LNP to induce cellular immunity. The GC responses are responsible for generating high-affinity neutralizing antibodies,^20^ while Tfh cells regulate the GC response.^21, 22^ Furthermore, memory CD4^+^ T cells and memory CD8^+^ T cells may provide long-term protection. In our study, GC B cells (defined as Fas^+^/GL7^+^ cells) and Tfh cells (defined as CXCR5^+^/PD-1^+^ cells) were evaluated in draining lymph nodes (DLNs) by flow cytometry 30 days after the first immunization. Flow cytometry results showed a significant increase in MPXV-specific GC B cells and Tfh cells from mRNA-A-LNP and mRNA-B-LNP-vaccinated mice compared with naive mice upon stimulation with MPXV-specific antigens (Fig. 3a-b). This suggests that mRNA-A-LNP and mRNA-B-LNP immunization can produce a long-lasting memory B cell effect. Then, specific CD4^+^ and CD8^+^ Tem cells (defined as CD44^+^/CD62L^-^ cells) in the spleen of immunised mice were further assessed. It was remarkable that mRNA-A-LNP and mRNA-B-LNP were more effective in induced MPXV-specific CD8^+^ Tem cells than CD4^+^ Tem cells (Fig. 3c-d). This demonstrated that our vaccine caused a memory T cell effect with the ability to kill virus-infected cells.

**Figure 3.**
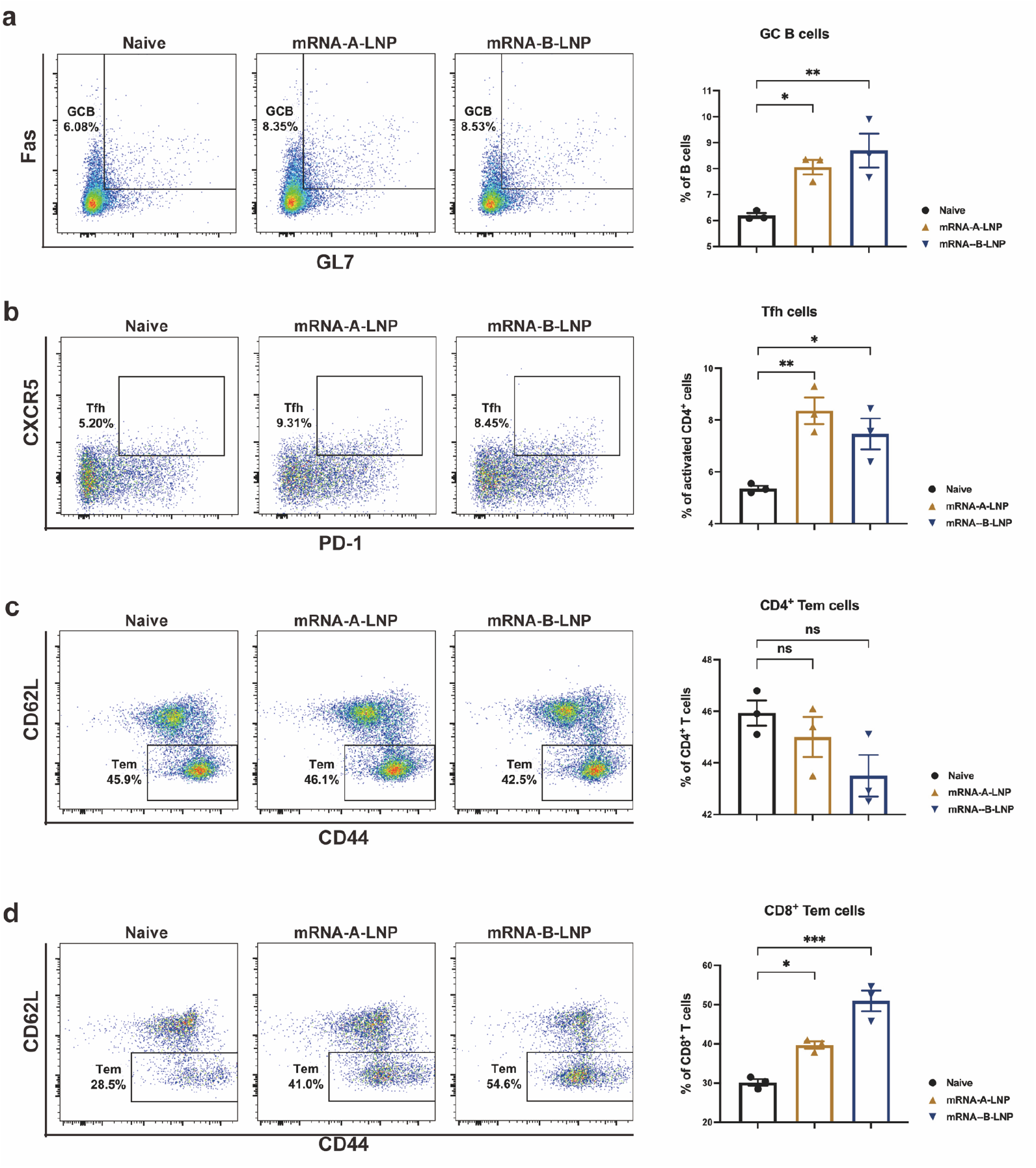
MPXV-Specific Cell Immune Response in mRNA-A-LNP and mRNA-B-LNP-Vaccinated Mice. (a-b) MPXV-specific GC B cells (a) and Tfh cells (b) in DLNs were detected by flow cytometry. (c-d) MPXV-specific CD4^+^ (c) and CD8^+^ (d) Tem cells in spleen were detected by flow cytometry. Data are shown as mean ± SEM. Significance was calculated using one-way ANOVA with multiple comparison tests (n.s., not significant; *p < 0.05, **p < 0.01, ***p < 0.001).

### The mRNA-A-LNP and mRNA-B-LNP protected mice from VACV challenge

The VACV challenge model based on firefly luciferase expression^23^ was used to assess the protection of mRNA-A-LNP and mRNA-B-LNP in mice. First, we investigated the passive protection of mRNA-A-LNP and mRNA-B-LNP immunized mouse sera (NT_50_: 6,284; 5,057), with sera from naive mice serving as negative controls. Viral infection in passively transferred serum nude mice was detected by bioluminescence imaging (BLI).^24^ At first, after 1 h of coincubation of serum with VACV *in vitro*, we observed a significant reduction in bioluminescent signal in 4-week-old nude mice injected with immunized mouse serum (Fig. 4a-b). It indicated that sera from mRNA-A-LNP and mRNA-B-LNP-immunized mice were effective in neutralizing VACV *in vitro*. In addition, the luminescence signal of 4-week-old nude mice was reduced by pre-injection of sera from immunized mice. Following the subcutaneous (s.c.) challenge 24 hours later, only a luminescence signal was detected at the injection site in nude mice passively immunized with mouse serum, indicating that the antibody had neutralized the virus (Fig. 4c-d). By contrast, significant viral infections were detected in nude mice passively transferred with serum from naive mice. These results demonstrated that mRNA-A-LNP and mRNA-B-LNP inoculated mouse sera had a passive protective efficacy against s.c. VACV challenge in immunodeficient young mice.

**Figure 4.**
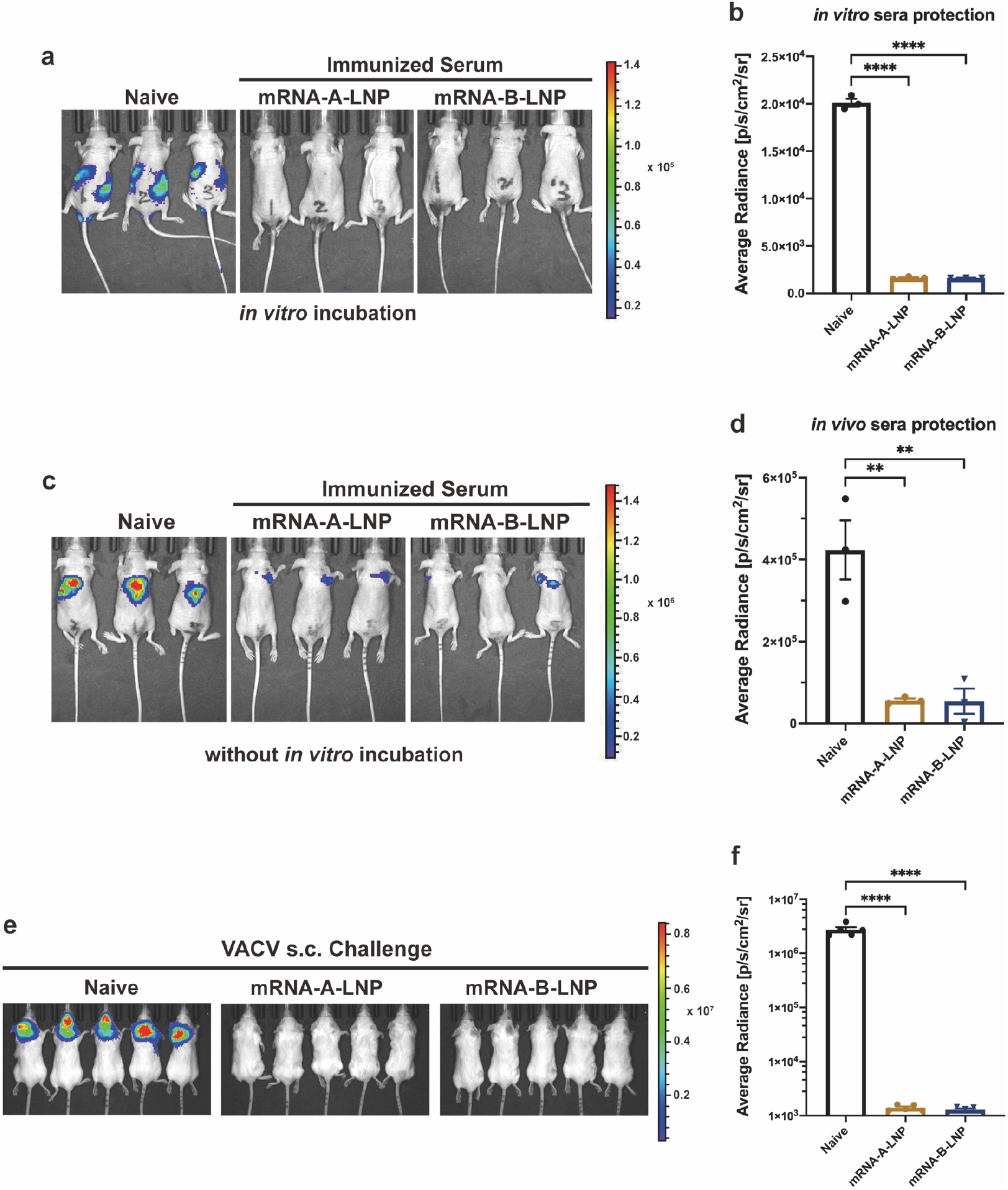
Protection of mRNA-A-LNP and mRNA-B-LNP against VACV Challenge in Mice. (a-b) *In vitro* serum protection. Serum (13 μl) and virus (4 × 10^3^ TCID_50_) were mixed for one hour before the i.v. and i.p. challenges of the 4-week-old nude mice (n = 3). Bioluminescent signal measurements were taken 6 hours after the viral challenge. (c-d) *In vivo* serum protection. Serum (50 μl) was first injected intravenously into the 4-week-old nude mice (n = 3), followed one hour later by s.c. challenge with VACV (2.5 × 10^5^ TCID_50_). Bioluminescent signal measurements were taken 24 hours after the viral challenge. (e-f) Thirty days after initial immunization, mice (n=5) were s.c. challenged with VACV (4 × 10^5^ TCID_50_). The viral load of mice was measured by bioluminescence imaging, and the bioluminescence signal of mice was measured 24 hours after the challenge. Data are shown as mean ± SEM. Significance was calculated using one-way ANOVA with multiple comparison tests (**p < 0.01, ****p < 0.0001).

Furthermore, we assessed the active protective efficacy of mRNA-A-LNP and mRNA-B-LNP. BALB/c mice immunized twice intramuscularly with 10 μg of mRNA-A-LNP or mRNA-B-LNP were challenged at day 30 with 4 × 10^5^ median tissue culture infectious doses (TCID_50_) of the VACV Tian Tan strain via the s.c. route. Viral load was measured in mice at 24 hours following the challenge by using BLI. The bioluminescent signal was largely undetectable in immune mice, whereas naive mice detected signal values as high as 6.5 log10, which suggested that VACV was rapidly cleared by vaccine-induced antibodies after the challenge (Fig. 4e-f). This suggests that prophylactic immunization with mRNA-A-LNP and mRNA-B-LNP is effective in protecting mice from s.c. VACV challenge.

### MPXV quadrivalent mRNA vaccine administered intramuscularly provides adequate safety

To assess the safety of our vaccine *in vivo*, we monitored several aspects, including mouse weight recordings, biochemical parameters and histopathological changes. After mRNA-A-LNP or mRNA-B-LNP immunization, body weight rapidly recovered, and no weight loss occurred (Fig. 5a). Since mRNA-A-LNP was prepared using the same method as mRNA-B-LNP, it was selected for further investigation of whether mice were harmed by the MPXV quadrivalent mRNA vaccine. First, according to clinical trial data, VACV-based vaccines have caused adverse events, including a 0.6% risk of myocarditis.^6^ Therefore, we further assessed the heart, liver and kidney function based on blood biochemical parameters. The data were within normal limits in the immunised and control groups, and there were no significant differences (Fig. 5b-f). Finally, multiple tissues from the mice were extracted for histopathological examination. As a result, there were no significant pathological changes between the immune and control groups (Fig. 5g). In conclusion, multiple lines of evidence demonstrate that our vaccine presents an adequate safety profile.

**Figure 5.**
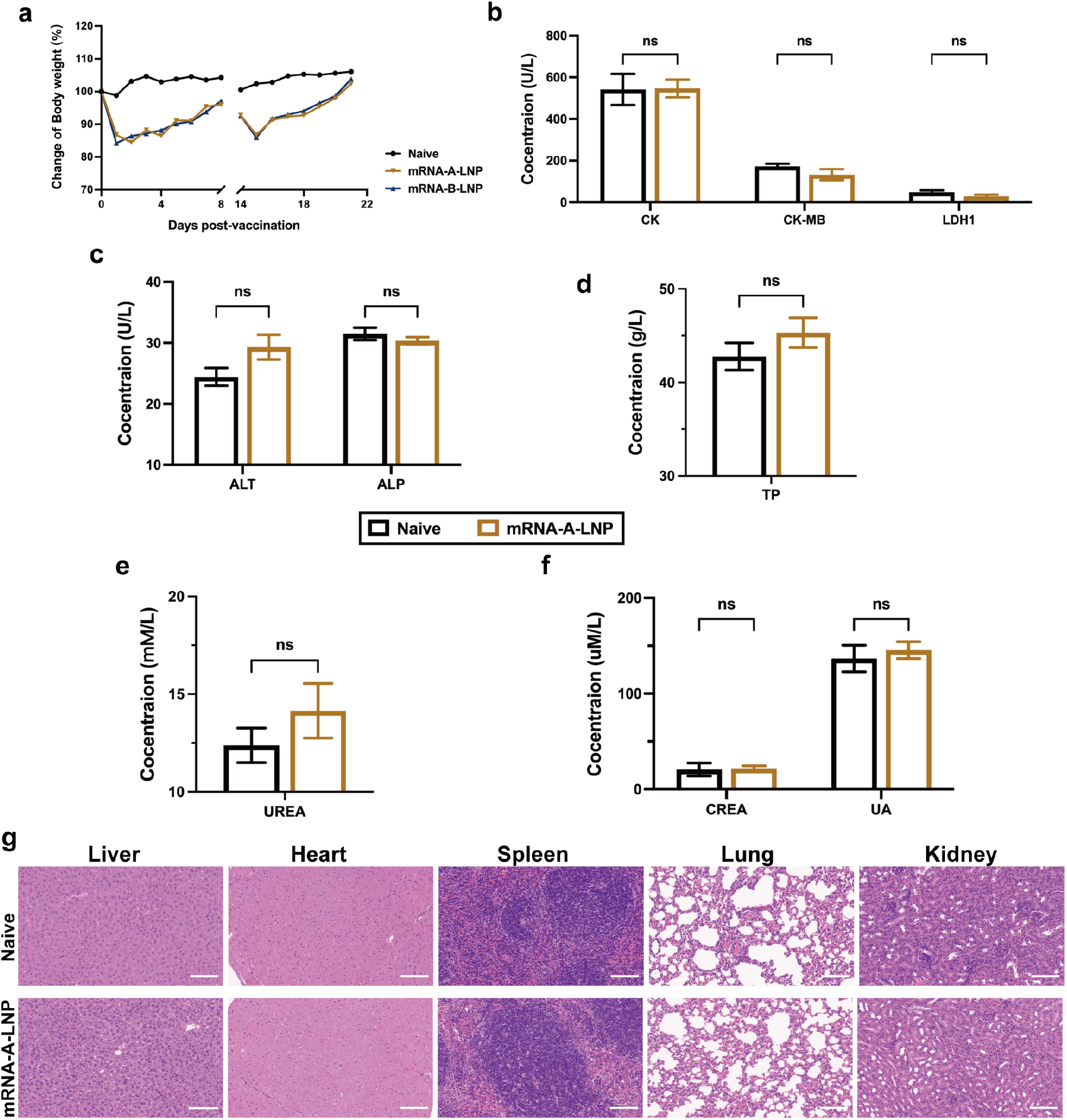
The safety evaluation of MPXV quadrivalent mRNA vaccine in mice. (a) The body weight records of mice on the first eight days after each vaccination. (b-f) Heart, liver and kidney function were determined by blood biochemical parameters (n = 5). CK, CK-MB and LDH1 represent heart function (b), ALT, ALP and TP represent liver function (c-d), while UREA, CREA and UA represent kidney function (e-f). (g) Representative histopathology (H&E) of different tissues, heart, liver, spleen, lung and kidney from naive mice or mRNA-A-LNP-immune mice. The H&E stained sections shown in the data are representative results from three test mice 48 hours post-inoculation. Scale bar = 100 μm, 30 ×. Data are shown as mean ± SEM. Significance was calculated using two-way ANOVA with multiple comparison tests (n.s., not significant; *p < 0.05, **p < 0.01).

## Discussion

Since the use of live VACV in the past century, smallpox has been completely eradicated, associated vaccination no longer occurs, and vaccine production has ceased accordingly. In this study, we report the immunogenicity and efficacy of the novel MPXV mRNA vaccine candidate, mRNA-A-LNP and mRNA-B-LNP. Two doses of immunization with mRNA-A-LNP or mRNA-B-LNP, respectively, induced potent MPXV IgG and robust VACV-neutralizing antibodies, and elicited durable MPXV-specific memory cellular immunity in mice.

The subunit-based smallpox vaccines have shown exciting feasibility in previous studies. Both combinatorial DNA vaccines and protein vaccines containing poxvirus antigens generated protective antibody responses in small animals and non-human primate models.^12, 15, 16, 25^ In our study, mRNA-A-LNP and mRNA-B-LNP vaccines produced high titers of IgG antibodies against the MPXV antigens A29L, A35R, M1R, and B6R (Fig. 2b-e). As MPXV A29L, A35R, M1R, and B6R genes exhibit high conservation with the orthologous genes of orthopoxviruses, including VACV and smallpox virus^16^, we speculate that mRNA-A-LNP and mRNA-B-LNP have cross-neutralizing effects on orthopoxviruses. Interestingly, live-virus neutralizing antibody assays determined that our MPXV vaccine elicited potent VACV-neutralizing antibodies (Fig. 2f). This means that MPXV vaccines provide cross-neutralizing antibodies against VACV within orthopoxviruses. Furthermore, vaccine-induced humoral responses were confirmed in nude mice by passive transfer tests.^26, 27^ It was observed that passive transfer of serum from vaccinated mice provides passive protection in nude mice against s.c. VACV challenge (Fig. 4a-d). This suggested that mRNA-A-LNP and mRNA-B-LNP-induced antibodies could provide protection in immunodeficient young mice. This is consistent with another study in which serum from DNA vaccinated animals were able to protect mice against a VACV challenge.^26^ Furthermore, two doses of prophylactic immunization with mRNA-A-LNP and mRNA-B-LNP protected mice from s.c. VACV challenge (Fig. 4e-f).

Using the mRNA vaccine platform, the MPXV quadrivalent mRNA vaccines, mRNA-A-LNP and mRNA-B-LNP, induced high titers of functional antibodies in mice while also boosting cellular immunity. The induction of cellular immunity may be a critical factor in explaining the remarkable protection provided by live attenuated vaccines compared with subunit vaccines.^12, 28^ In the application of the SARS-CoV-2 vaccine, mRNA vaccines have been well demonstrated to induce cellular immunity.^20, 29^ It is consistent with our findings that the MPXV quadrivalent mRNA vaccine was able to elicit excellent cellular immunity (Fig. 3a-d). In addition, live virus vaccines do not stop the virus from spreading.^30^ In contrast, the smallpox DNA vaccine, a nucleic acid vaccine, has been reported to prevent the shedding of infectious viruses in the oral cavity of vaccinated animals.^31^ As a result, it is reasonable to hypothesize that the MPXV mRNA vaccine, also as a nucleic acid vaccine, will prevent the shedding of infectious viruses in all inoculated animals. And we will conduct further studies to test this hypothesis.

In the orthopoxvirus route of infection, monocytes are first recruited to the site of infection as the initial target.^32, 33^ The cytolytic T cells, meanwhile, kill infected monocytes to prevent virus spread. Memory CD8^+^ T cells have been reported effective in protecting susceptible mice from lethal orthopoxvirus challenges.^34-36^ Since mRNA-A-LNP and mRNA-B-LNP primarily elicit memory CD8^+^ T cells, allowing T cell-mediated lysis during the early infection period of viral challenge would prevent virus spread (Fig. 3d). Furthermore, circulating antibodies performed a similar role to CD8^+^ T cells.^34^ It is the long-lived plasma cells (LLPC) and memory B cells that are responsible for most of the prolonged humoral immunity induced by vaccines.^37^ Memory B cells induced with the smallpox vaccine could respond quickly to infection and replenish LLPC to maintain long-term antibodies levels in humans, according to a study.^38^ Moreover, memory B cells were also associated with IgM - IgG isotype switching. Therefore, considering that memory B cells are developed in the germinal center, the significant increase in MPXV-specific GC B cells and Tfh cells suggests that mRNA-A-LNP and mRNA-B-LNP can maintain protective antibody responses with high affinity and durability (Fig. 3a-b). In summary, we report efficient and safe quadrivalent mRNA vaccine candidates against MPXV, based on MPXV-specific antigens A29L, A35R, M1R and B6R. The vaccines reported here are the first MPXV vaccines developed using an mRNA vaccine platform. As mRNA-A-LNP and mRNA-B-LNP induce solid humoral and cellular immunity, they could provide new ideas for orthopoxvirus vaccine development. Considering the rapidly spreading monkeypox epidemics, MPXV mRNA vaccines which can be rapidly developed and prepared on a large scale, are expected to protect against infection-related symptoms, hospitalizations, and death.

## Materials and methods

### Ethics statement

All animal studies, there were reviewed and approved by the Animal Experiment Committee of Laboratory Animal Center, Academy of Military Medical Sciences (AMMS), China (Assurance Number: IACUC-DWZX-2022-576). All animal studies were conducted strictly in accordance with the guidelines set by the Chinese Regulations of Laboratory Animals and Laboratory Animal Requirements of Environment and Housing Facilities.

### Cells and viruses

HEK293T, Huh-7, RD, Vero and 143TK cells were cultured in Dulbecco’s Modified Eagle Medium (DMEM; Thermo Fisher) supplemented with 10 % fetal bovine serum (FBS; Thermo Fisher) and penicillin (100 U/ml)-streptomycin (100 mg/ml) (Thermo Fisher). All cells were grown at 37 °C in a humidified 5% CO_2_ atmosphere.

The vaccinia virus (VACV) Tian Tan strain expressing firefly luciferase was propagated in 143TK cells and titrated in Vero cells.^23^

### Synthesis and characterization of MPXV mRNA

All mRNA sequences encoding MPXV proteins (A29L, A35R, M1R, B6R) were prepared by *in vitro* transcription, as described previously.^22^ Transcription was performed from linearized DNA templates using the T7-FlashScribe™ Transcription Kit (Cellscript). Further, the modified mRNA was synthesized by replacing UTP in the kit with pseudo-UTP (TriLink).

Afterwards, the RNA was capped using the ScriptCap™ Cap 1 Capping System kit with ScriptCap™ Capping enzyme and 2’-O-methyltransferase (Cellscript) according to the manufacturer’s instructions. The mRNA product purified by ammonium acetate precipitation was then resuspended in RNase-free water for further analysis and application.

The concentration and quality of the synthesized MPXV mRNA were measured using an Agilent 2100 Bioanalyzer and RNA Nano 6000 Assay Kit (Agilent), according to the manufacturer’s instructions.

All mRNAs (2 μg) were transfected into HEK293T cells using Lipofectamine 3000 transfection reagent (Thermo Fisher) according to the manufacturer’s instructions, lysates were collected, and Western blotting was performed.

### Formulation and characterization of mRNA-LNP

Lipid nanoparticle (LNP) formulations were prepared using NanoAssemblr Ignite’s (Precision Nanosystems) NxGen Microfluidics technology. Briefly, lipids containing ionised lipids, 1, 2-distearoyl-sn-glycero-3-phosphocholine (DSPC), cholesterol and DMG-PEG2000 were dissolved in ethanol (with a molar ratio of 50:10:38.5:1.5). In an Ignite™ mixer, the lipid mixture was combined with 20 mM citrate buffer (pH 4.0) containing mRNA in a 1:3 volume ratio. It was then diluted through a 100 k MWCO PES membrane (Sartorius Stedim Biotech) against a 10-fold volume of DPBS (pH 7.4) and concentrated to the desired concentration.

The particle size and ζ-potential of mRNA-LNP were measured by a Litesizer 500 (Anton Paar). Particle size measurements were carried out using dynamic light scattering (DLS). The data were also analyzed using the Anton Paar Kalliope software package.

The morphology of mRNA-LNP was analyzed by transmission electron microscopy (Hitachi H-7800, Tokyo) using a negative staining technique. mRNA-LNP was absorbed into the copper mesh for 60 s and stained with phosphotungstic acid (1%) for 20 s before observation.

The Quant-iT RiboGreen RNA Reagent and Kit were used to detect the encapsulation of mRNA-LNP according to the manufacturer’s instructions

### Mouse vaccination

In this experiment, 15 BALB/c mice (6-8 weeks of age, female, SPF) were randomly divided into three groups (n = 5). The mice were immunized with the mRNA-A-LNP vaccine or the mRNA-B-LNP vaccine or were designated negative controls. We administered the vaccine intramuscularly at 40 μg on day 0. And a booster immunization was administered on day 14. All mice were tested for IgG and neutralizing antibodies on days 10 and 24 after the initial immunization. The following flow cytometry analyses were performed 30 days after the initial vaccination.

### Evaluation of serum antibody

IgG antibody titers to MPXV-specific antigens A29L, A35R, M1R and B6R were determined by enzyme-linked immunosorbent assay (ELISA). VACV-specific neutralizing antibody titers were determined by a live-virus based neutralization test.

#### (a) ELISA assay

IgG antibody titers against MPXV-specific antigens A29L, A35R, M1R and B6R were determined by ELISA. A 96-well plate was coated with 1 μg/ml of A29L (TSP-MV002, Tsingke Biotechnology), A35R (TSP-MV003, Tsingke Biotechnology), M1R (TSP-MV005, Tsingke Biotechnology) or B6R (40902-V08H, Sino Biological) protein respectively and incubated overnight at 4°C.

After incubation, plates were washed with 1 x TBST and blocked with BSA for 2 hours at 37 °C. Then, serial 2-fold gradient dilutions of serum starting at 1:100 were added to the wells, diluted with casein block, and incubated for 1 hour at 37 °C. Next, the plates were washed and treated with horseradish peroxidase (HRP)-conjugated goat anti-mouse IgG (Abclonal) for 1 hour at 37 °C. Afterwards, the plates were washed and incubated with the substrate tetramethylbenzidine (TMB; TIANGEN) for 20 minutes at room temperature in the dark before the reaction was terminated with hydrochloric acid (2M; Solarbio). Absorbance at 450/630 nm was recorded using an I-control Infinite 200 PRO microplate reader (TECAN). The ELISA endpoint titers were defined as the dilution of vaccinated serum, which resulted in absorbance no less than 2.1-fold that of the average negative serum (1:100).

#### (b) Live-virus neutralization assay

Neutralizing antibody titers were determined by a live-virus neutralization assay. The VACV Tian Tan strain was designed to encode firefly luciferase protein.^39^ Sera were tested for neutralizing activity against the live virus via mixing serial 3-fold diluted sample, starting at 1:30, with 4 × 10^3^ TCID_50_ of VACV. The serially diluted serum samples were mixed with the diluted virus in an equal volume. The antibody–virus and virus-only mixtures were then incubated at 37 °C with 5% CO_2_. After incubating for 1 hour, we added Vero cells (40,000 cells/well) to each well at 37 °C with 5% CO_2_. After 48-hour incubation, the cells were lysed, and the luciferase activity was measured via Bright-Glo Luciferase Assay System (Promega) according to the manufacturer’s specifications. Luciferase activity was then measured using an EnSight plate reader (PerkinElmer). Neutralizing activity was calculated by quantification of luciferase activity in relative light units (RLU). 50% live-virus neutralizing antibody titer (NT_50_) were calculated using a log (inhibitor) vs. normalized response (Variable slope) non-linear regression model in GraphPad Prism 8.0 (GraphPad Software).

### Evaluation of cellular immune response

Cells from spleen and DLNs were isolated and analyzed by flow cytometry to determine the cellular immune response. DLNs were used to analyze Tfh cells responses and GC B cells responses, Spleen was used to analyze CD4^+^ or CD8^+^ Tem cells responses.

Briefly, cells from spleen or DLNs (1,000,000 cells/well) were stimulated with MPXV-specific antigens A29L, A35R, M1R and B6R (2 μg/ml each protein) at 37 °C in 5% CO_2_ for 12 h. Brefeldin A (5 μg/ml; Biolegend) was incubated with cells for 4 h. Then, Fc receptors of cells were blocked using CD16/CD32 antibodies (Mouse BD Fc Block; BD Biosciences) for 15 min at 4 °C, and cells were stained with a cocktail of fluorescently conjugated antibodies to CD3-PE/Cyanine7 (Biolegend), CD4-FITC (Biolegend), CD8-PercP (Biolegend), CD44-PE (Biolegend), CD62L-APC (Biolegend), B220-FITC (Biolegend), CD4-PercP/Cyanine5.5 (Biolegend), CD44-APC (Biolegend), PD-1-PE/Cyanine7 (Biolegend), B220-PercP/Cyanine5.5 (Biolegend), Fas-PE (Biolegend) and GL7-FITC (Biolegend) for another 30 min at 4 °C in dark. Following washing with cell staining buffer (BD Biosciences), dead cells were stained with Fixable Viability Dye eFluor™ 780 (Thermo Fisher Scientific) for 30 min at 4 °C in the dark. A final wash with cell staining buffer, data were obtained by FACS Aria II flow cytometer (BD Biosciences) and analyzed by Flow J software. The GC B cell response was represented as live^+^/B220^+^/Fas^+^/GL7^+^. The Tfh cell response was represented as live^+^/B220^-^ /CD4^+^/CD44^+^/PD-1^+^/CXCR5^+^. The CD4^+^ Tem cell response was represented as live^+^/CD3^+^/CD4^+^/CD44^+^/CD62L^-^. The CD8^+^ Tem cell response was represented as live^+^/CD3^+^/CD8^+^/CD44^+^/CD62L^-^.

### Serum protective test

The VACV challenge model is based on the VACV Tian Tan strain expressing firefly luciferase. Using 4-week-old BALB/c nude mice, the serum passive protection model was developed. Protective sera were collected from BALB/c mice immunised with mRNA-A-LNP and mRNA-B-LNP. Serum samples were collected 24 days after the initial immunization. To assess *in vitro* sera protection, serum (13 μl) and virus (4 × 10^3^ TCID_50_) were mixed for one hour before intravenous (i.v.) and intraperitoneal (i.p.) challenge of nude mice. To assess *in vivo* sera protection, serum (50 μl) was first injected intravenously into nude mice, followed one hour later by s.c. challenge with VACV (2.5 × 10^5^ TCID_50_). Bioluminescent signal measurements were made following the viral challenge.

Bioluminescence imaging (BLI) was acquired and analyzed using the IVIS Lumina Series III imaging system (PerkinElmer). Briefly, luminescence was measured 5 minutes after intraperitoneal injection of the substrate D-luciferin (PerkinElmer). The bioluminescent signals in regions of interest (ROIs) were quantified using Living Image 3.5.

### Mouse challenge

The VACV challenge model is based on the VACV Tian Tan strain expressing firefly luciferase. BALB/c mice (n=5) immunized with mRNA-A-LNP and mRNA-B-LNP were challenged subcutaneously with VACV (4 × 10^5^ TCID_50_) 30 days after the initial immunization. At hour 24 post-challenge, mice were measured for bioluminescence signals.

### *In vivo* toxicity

To assess the *in vivo* toxicity of the vaccine, body weights were recorded after vaccination. Mice vaccinated with mRNA-A-LNP (40 μg; n = 3) were analyzed for heart, liver and kidney function 48 hours after immunisation using a Chemray 240 and Chemray 800 (Rayto) automated biochemical analyzer.

Organ tissues, including heart, liver, spleen, lung and kidney, were extracted 48 hours after injection for histopathology and fixed in 4% neutral buffered formaldehyde. Afterwards, they were embedded in paraffin, sectioned and stained with hematoxylin and eosin (H&E). Images were taken with a NIKON Eclipse CI microscope.

### Statistical analysis

Statistical analyses were performed using GraphPad Prism 8.0 (GraphPad Software). All of the data are presented as the mean ± SEM. Statistical difference was analyzed by one-way or two-way ANOVA. All tests are accepted as statistically significant when the *p* value is less than 0.05.

## ACKNOWLEDGEMENTS

The study was supported by the National Key R&D Program of China (2021YFC2302405) and the National Natural Science Foundation of China (Grant No. 81830101).

## AUTHOR CONTRIBUTIONS

JY and SW conceived the project. YS and ZZ synthesized the mRNA vaccine and performed the experiments. FL, YW and WH performed animal challenge experiments. HL, CY, HS, JL, YC, JM, XW, and JF provided experimental support. JY designed the MPXV mRNA sequence. YS, ZZ and JY analyzed all the data and wrote the manuscript. JY and SW edited and revised the manuscript. All authors read and approved the final version of the manuscript.

